# Environmental enrichment mitigates stroke-induced change in sharp-wave associated ripple characteristics

**DOI:** 10.1101/2021.02.12.431002

**Authors:** Zachary Ip, Gratianne Rabiller, Ji-Wei He, Shivalika Chavan, Yasuo Nishijima, Yosuke Akamatsu, Jialing Liu, Azadeh Yazdan-Shahmorad

## Abstract

Cognitive and memory impairments are common sequelae after stroke, yet how middle cerebral artery (MCA) stroke chronically affects the neural activity of the hippocampus, a brain region critical for memory but remote from the stroke epicenter, is poorly understood. Environmental enrichment (EE) improves cognition following stroke; however, the electrophysiology that underlies this behavioral intervention is still elusive. We recorded local field potentials simultaneously from sensorimotor cortex and hippocampus in rats following MCA occlusion and subsequent EE treatment. We found that MCA stroke significantly impacted the electrophysiology in the hippocampus, in particular it disrupted characteristics of sharp-wave associated ripples (SPW-Rs) which are known correlates of memory and cognition. Importantly, we show that EE mitigates stroke-induced changes to SPW-R characteristics. These results begin to uncover the complex interaction between cognitive deficit following stroke and EE treatment, providing a testbed to assess different strategies for therapeutics following stroke.

## Introduction

Stroke is a leading cause of adult disability, with the most common occurrence in the middle cerebral artery (MCA) region in humans. Unfortunately, there are few effective treatment options for disability following stroke besides physical therapy. In addition to the impairment of high-level sensorimotor functions, a common outcome of stroke is cognitive and memory deficit (Khedr 2009, Cumming 2009, Barker-Collo 2010). The hippocampus is highly involved in the encoding and retrieval of memories, but hippocampal and parahippocampal areas are rarely directly affected by MCA stroke because hippocampal blood flow is supplied by the posterior circulation (Bederson 1986, Iizuka 2011, Liu and McCullough 2011). Animal models represent this phenomenon well, displaying cognitive impairment following stroke in the absence of hippocampal injury (Okada 1995, Wang 2011, Sun 2013). However, a thorough understanding of the mechanisms underlying cognitive and memory impairment caused by MCA stroke remains poorly understood.

Cortical dysfunction following MCA stroke has been well described using both histological and electrophysiological methods (Oliveira 2014, Hazime 2020). Hippocampal functional impairment following MCA stroke has been demonstrated using behavioral assessment (Wang 2011, Barth 2011), however how hippocampal electrophysiology changes following cortical lesioning has not been studied. There are many hippocampal electrophysiological features that can be used to interrogate memory function such as the ratio of theta band to delta band signal power (TD) of the CA1 pyramidal layer within the hippocampus which defines brain states relevant to memory function (Ognjanovski 2014, Aminov 2017). High theta/delta ratio (HTD) is correlated with memory performance and memory consolidation during rapid eye movement sleep (Buzsaki 2002, Battaglia et al 2011). Manipulation of hippocampal HTD alters cognition, further supporting HTD’s role in cognition (Williams and Tortella 2002, Aminov 2017). Meanwhile, low theta/delta ratio (LTD), also known as slow wave state, has been associated with immobility, during which the hippocampus experiences sharp-wave associated ripples (SPW-Rs) (Kay 2016). SPW-Rs are short, high frequency oscillations within the CA1 pyramidal layer of the hippocampus that are concurrent with a negative deflection in the radiatum layer that represent memory recall and encoding (Carr 2011; Jadhav 2012; Buzsaki 2015).

Theta oscillations in the hippocampus are known to provide a temporal reference for local computations by modulating high frequency gamma oscillations in what is known as theta-gamma coupling (Lisman 2008; Hanslmayr 2016, Heusser 2016) which can be measured using cross frequency phase amplitude coupling (PAC) (Tort 2010). Theta-gamma coupling within the hippocampus has been shown to support memory processes and occurs during HTD (Tort 2009; Shirvalkar 2010; Colgin 2015). It has also been observed between brain regions such as the prefrontal and entorhinal cortex (Tamura 2017; Bandarabadi 2019). Manipulation of theta rhythms in the hippocampus alters cognitive performance (McNaughton 2006), further supporting theta oscillations causative role in cognition.

Chronic stroke leads to a complex cascade of effects within the brain such as the loss of functional connectivity (Silas and Murphy 2014, Schmitt et al 2017) and changes in local oscillations (Rabiller et al 2015, Zhang et al 2006, Moyanova and Dijkhuizen 2014, Ip et al 2019) which have the potential to effect remote brain areas such as the hippocampus. EE is an effective non-invasive therapy that has long been studied as a potential treatment for improving cognition (Cooper and Zubek 1958; Diamond 1966; Manosevitz 1970; Nilsson 1999), by increasing exposure to novelty, social contact, and physical activity. Cognitive and behavioral deficit following stroke has consistently been shown to be improved by environmental enrichment (EE) (Hamm 1996; Passineau 2001; Ip 2002; Komitova 2005; Matsumori 2006; Fan 2007; Wang 2011; Wang 2019). However, the underlying electrophysiological mechanisms are still largely unknown.

Our previous work has shown that an acute reduction in cerebral blood flow caused by MCA occlusion disrupts the electrophysiology of the hippocampus, which we observed through aberrant increases in SPW-R frequency and theta-gamma coupling between hippocampus and cortex within the first hour of ischemia (He 2019). Here we seek to understand the changes in hippocampal electrophysiology during chronic phase of MCA stroke to interrogate the underlying mechanisms of cognitive impairment following stroke and cognitive improvement following EE.

## Results

After inducing a lesion by unilateral distal middle cerebral artery occlusion (dMCAO), we randomly assigned rats to standard and EE housing groups during recovery. We divided the animals into two time-point stroke subgroups and a non-stroke control group for recording under urethane anesthesia before sacrifice. All groups were composed of different animals: control (n = 8), 2 weeks post-stroke (2WS) (n = 7), and 1 month post-stroke (1MS) (n = 10). The EE group was split into EE control (EEC) (n = 10), and 1-month post stroke (EES) (n = 9).

We analyzed both the absolute infarcted volume as well as the ratio of infarcted volume to intact tissue volume to confirm there was no hippocampal lesion and determine whether lesion size was affected by the chronicity of stroke or by exposure to enrichment. There was no apparent morphological difference in the hippocampus as revealed by hematoxylin and eosin (H&E) staining between the stroke and non-stroke groups, suggesting that distal occlusion of MCA did not compromise hippocampal structural integrity. Both analyses revealed that lesion size did not significantly differ between groups (ANOVA; absolute p = 0.460, ratio p = 0.485) (Supplemental Figure 1). The locations of probes were verified by histology, spanning from −3mm to −3.72mm AP and 2.5mm to 3mm laterally (Figure 1).

**Figure 1.**
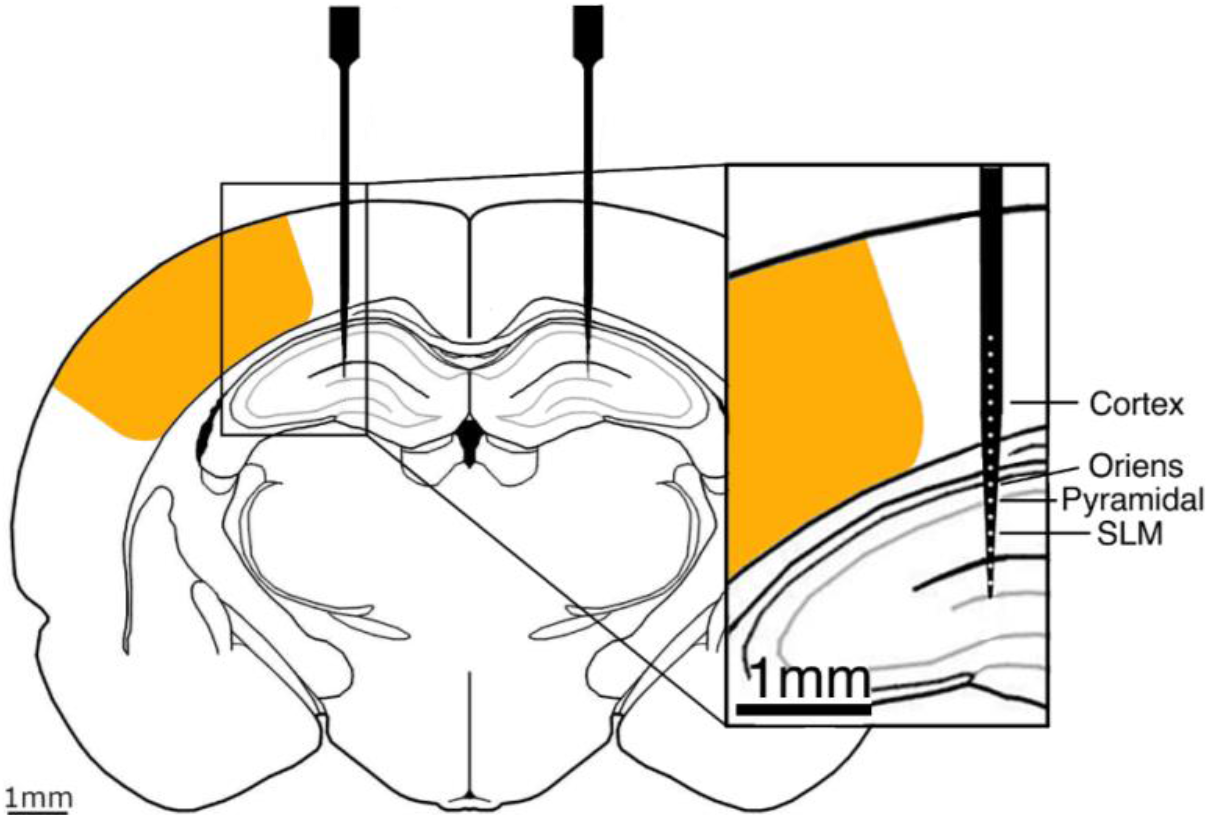
Schematic of infarct area and probe locations. Probes are inserted to cover sensorimotor cortex and hippocampus. Approximate infarct and peri-infarct areas from stroke indicated by orange shading.

We analyzed normalized signal power within cortex and hippocampus as a simple metric of activity levels within the tissue. Surprisingly, there were very sparse significant changes between groups. Delta signal power of 2WS and 1MS tended to be lower than control in both cortex and hippocampus, while interestingly theta, gamma and high gamma signal power tended to be higher in both 2WS and 1MS compared to control (Supplementary Figure 2).

### Brain state stability is disrupted following stroke

Under urethane anesthesia, the brain experiences sleep-like activity. The ratio of theta band to delta band signal power in the pyramidal layer of the hippocampus during sleep defines states relevant to memory. We analyzed the stability of theta/delta brain state under anesthesia by analyzing the duration of HTD state (Figure 2A). Surprisingly, we found that TD state stability is disrupted following stroke despite no direct lesion to the hippocampus, with a significant decrease in the duration of HTD brain state bilaterally for both stroke groups compared to control (ANOVA; ipsilesional; 2WS p = 4.86e-7, 1MS p = 3.33e-4, contralesional; 2WS p = 1.41e-9, 1MS p = 5.13e-7) (Figure 2B, Supplementary Figure 3A). However, the disruption of state stability does not alter the overall proportion of HTD to LTD as evidenced by the HTD/LTD ratio (Figure 2C, Supplementary Figure 3B) (ANOVA; p > 0.21). This shows that stroke chronically disrupts the stability of brain states defined within the hippocampus, but does not disrupt the proportion of HTD to LTD.

**Figure 2.**
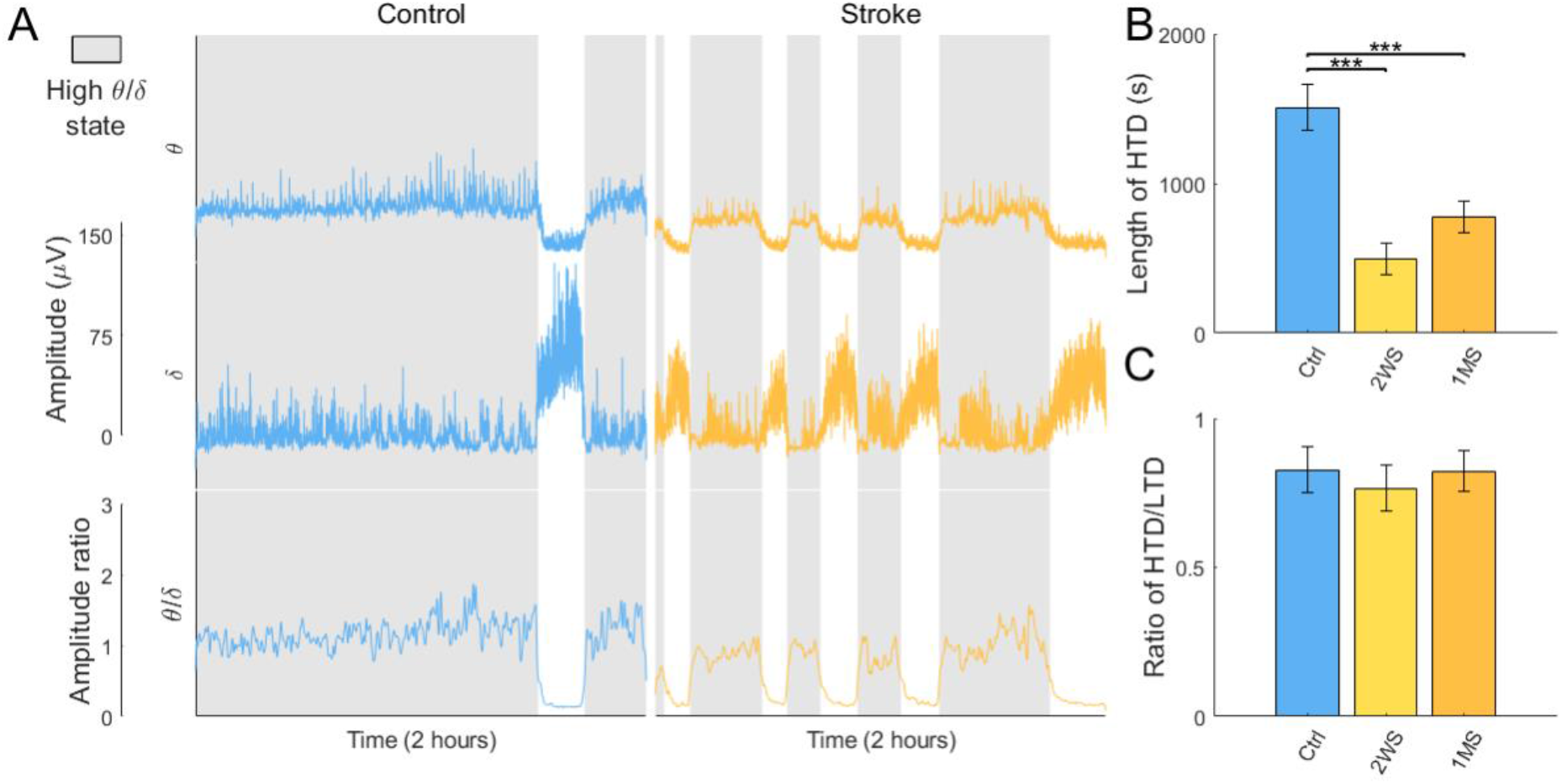
Detecting HTD and LTD states. (A) The columns show examples of LFP traces for a control and stroke sample. The rows are: 1-Spontaneous amplitude of theta, 2-Spontaneous amplitude of delta, 3-Ratio of theta/delta. (B) Comparison of the average duration of ipsilateral HTD state. (C) Comparison of the proportion of ipsilesional HTD to LTD. *Significant differences (p < 0.001, are demarked with ***).*

### SPW-R characteristics change following stroke

SPW-Rs occur within the CA1 pyramidal layer of the hippocampus during LTD and represent memory encoding. To quantify affected memory performance, we analyzed characteristics of SPW-Rs. There was an increase in SPW-R signal power of both ipsilesional and contralesional hemispheres in 2WS and 1MS compared to control (Kruskal Wallis; ipsilesional; 2WS: p = 9.97e-4, 1MS: p < 1e-16 contralesional 2WS p = 3.00e-16, 1MS p < 1e-16) (Figure 3A, Supplementary Figure 4A). The duration of SPW-Rs was also significantly different; SPW-Rs at 2WS were significantly longer than control (ANOVA; p = 5.41e-5), while SPW-Rs at 1MS were significantly shorter (ANOVA; p = 5.41e-5) (Figure 3B). These results indicate that stroke significantly affects both power and duration of SPW-Rs. Both the signal power and duration of SPW-Rs at 2WS significantly increased compared to 1MS in both hemispheres (Kruskal Wallis; signal power; ipsilesional p < 1e-16, contralesional p < 1e-16, duration; ipsilesional p = 4.03e-9, contralesional p < 1e-16). These results indicate that there is some compensatory mechanism occurring at 2WS.

**Figure 3.**
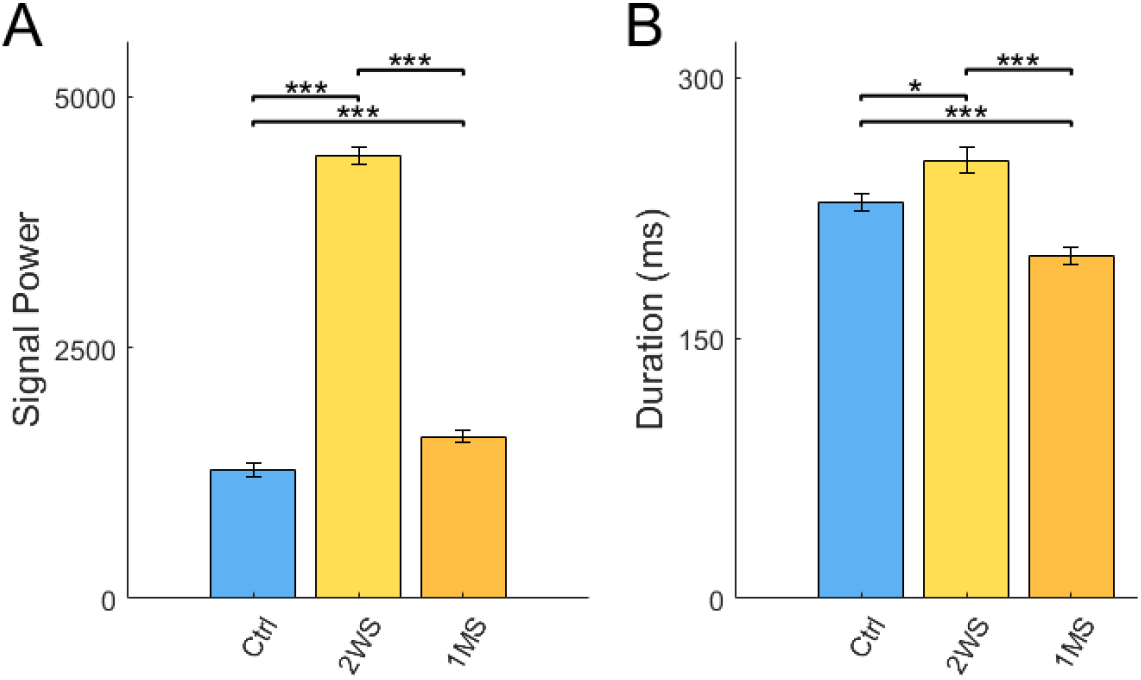
Comparison of ipsilesional (A) SPW-R power and (B) SPW-R duration. *Significant differences (p < 0.05, p < 0.01, andp < 0.001 are demarked with *, **, or *** respectively).*

### Current flow surrounding SPW-Rs is disrupted following stroke

We performed laminar current source density analysis (CSD) aligned to the onset of SPW-R to evaluate current flow through the hippocampus during SPW-Rs. The control group revealed pairs of dipoles with the apparent source centered in the pyramidal layer with the sink centered in the radiatum as expected. After SPW-R, the dipole reverses at a lower amplitude, with the sink in pyramidal and the source in the radiatum (Figure 4A). This post-SPW-R phase lasts approximately 0.6 seconds before dissipating. To analyze changes to this current flow pattern, we defined windows of interest before, during, and after SPW-R to evaluate the strength of related current flow.

**Figure 4.**
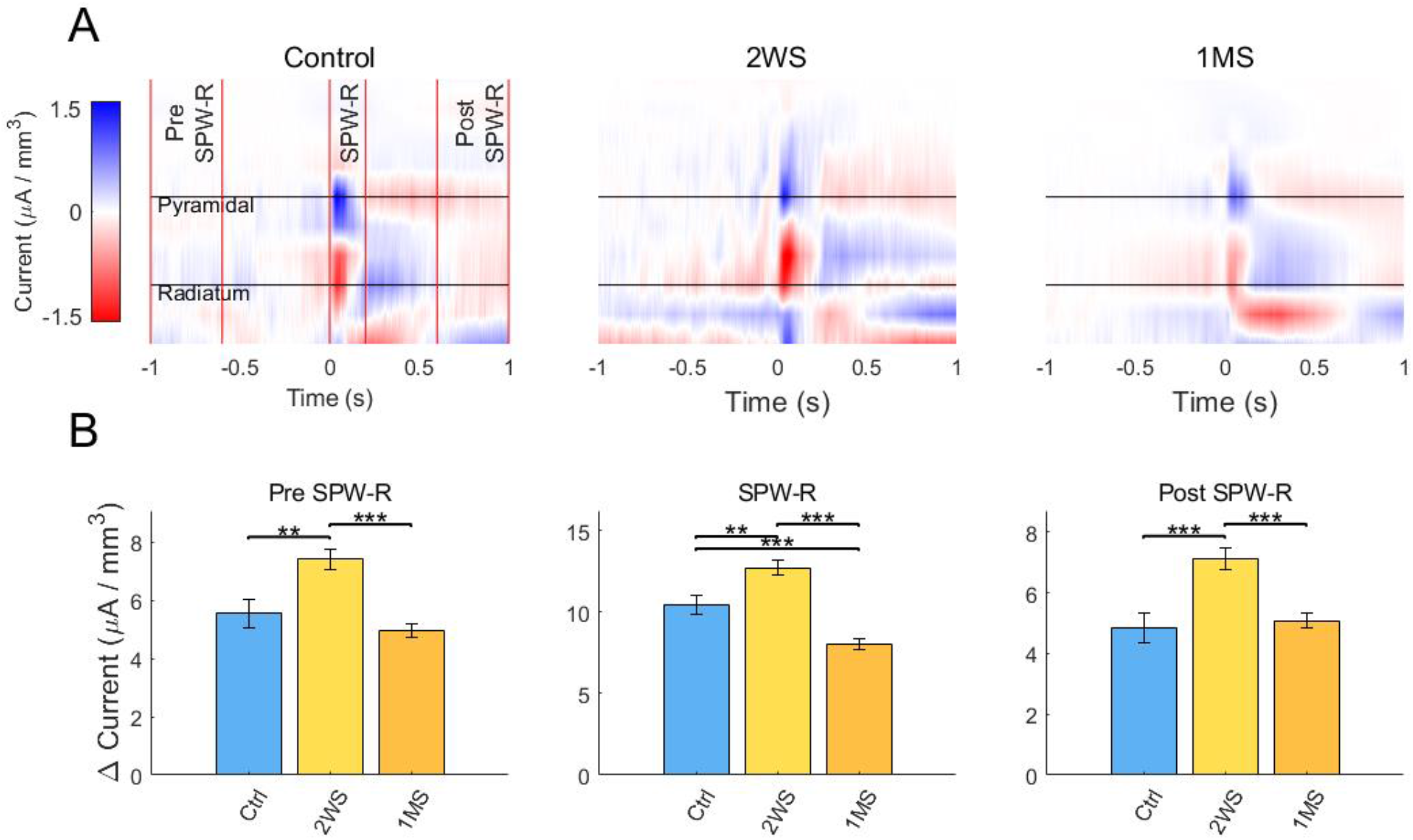
Comparison of ipsilesional CSD during and following SPW-R. (A) CSD plots of ipsilesional hemisphere, displaying average current of all ripples for all animals within a particular group. Windows of interest are demarked with red lines. (B) Change in current was measured using difference between the minimum and maximum amplitudes with the demarked windows. *Significant differences: p < 0.01 and p < 0.001, are demarked with **, or *** respectively.*

In both hemispheres, the dipole amplitude before, during, and after SPW-R was significantly higher at 2WS compared to control (ANOVA; before; ipsilesional p = 0.0074, contralesional p = 1.24e-4, during; ipsilesional p = 0.0077, contralesional p = 0.0011, after; ipsilesional p = 5.83e-4, contralesional p = 0.0056) (Figure 4B, Supplementary Figure 5) while the amplitude of 1MS is significantly lower compared to control (ANOVA; ipsilesional p = 6.78e-4, contralestional 1.03e-5) (Figure 4B). Like SPW-R power and duration, the dipole amplitude at 1MS is significantly lower than 2WS before, during and after SPW-R (ANOVA; before; ipsilesional p = 1.29e-7, contralesional p = 6.32e-13, during; ipsilesional p < 1e-16, contralesional p < 1e-16, after; ipsilesional p = 1.11e-5, contralesional p = 4.72e-5). These results show that stroke causes significant change in current flow, while the decrease of dipole amplitude from 2WS to 1MS support our SPW-R previous observations that there is some compensatory activity at 2WS.

### Theta-gamma coupling between hippocampus and cortex is reduced following stroke

Theta rhythms coordinate high frequency gamma activity within the hippocampus during HTD and supports memory processes. We used PAC to detect theta-gamma coupling within the hippocampus, and to determine whether coupling existed between cortex and hippocampus. During HTD theta-gamma coupling and delta-high gamma coupling was present bi-directionally within the hippocampus as expected, however we also detected coupling between cortex and hippocampus in the control group. Coupling within the pyramidal layer and between pyramidal theta and cortical gamma are shown as examples (Figure 5A). Ipsilesional coupling within the pyramidal layer was significantly lower at 1MS compared to control. Interestingly, ipsilesional coupling between hippocampus and cortex at 1MS was also significantly lower than control for all hippocampal layers in compared to control (Figure 5B) (ANOVA; pyramidal p = 0.0085, SLM p = 0.00107, oriens p = 0.0356). Coupling within the hippocampus and between cortex and hippocampus is lower than control at 2WS, though not significantly. This could be due in part to the compensatory mechanisms observed in SPW-Rs. During LTD, theta-gamma coupling was not present within the cortex or between cortex and hippocampus as expected. Instead, only delta-high gamma coupling was present during LTD, which did not change following stroke. The breakdown in PAC between hippocampal theta and cortical gamma implies that MCA stroke, which does not cause infarct to the hippocampus, breaks down coordination of oscillations between theta and gamma within the hippocampus, and the coordination of cortical gamma by hippocampal theta.

**Figure 5.**
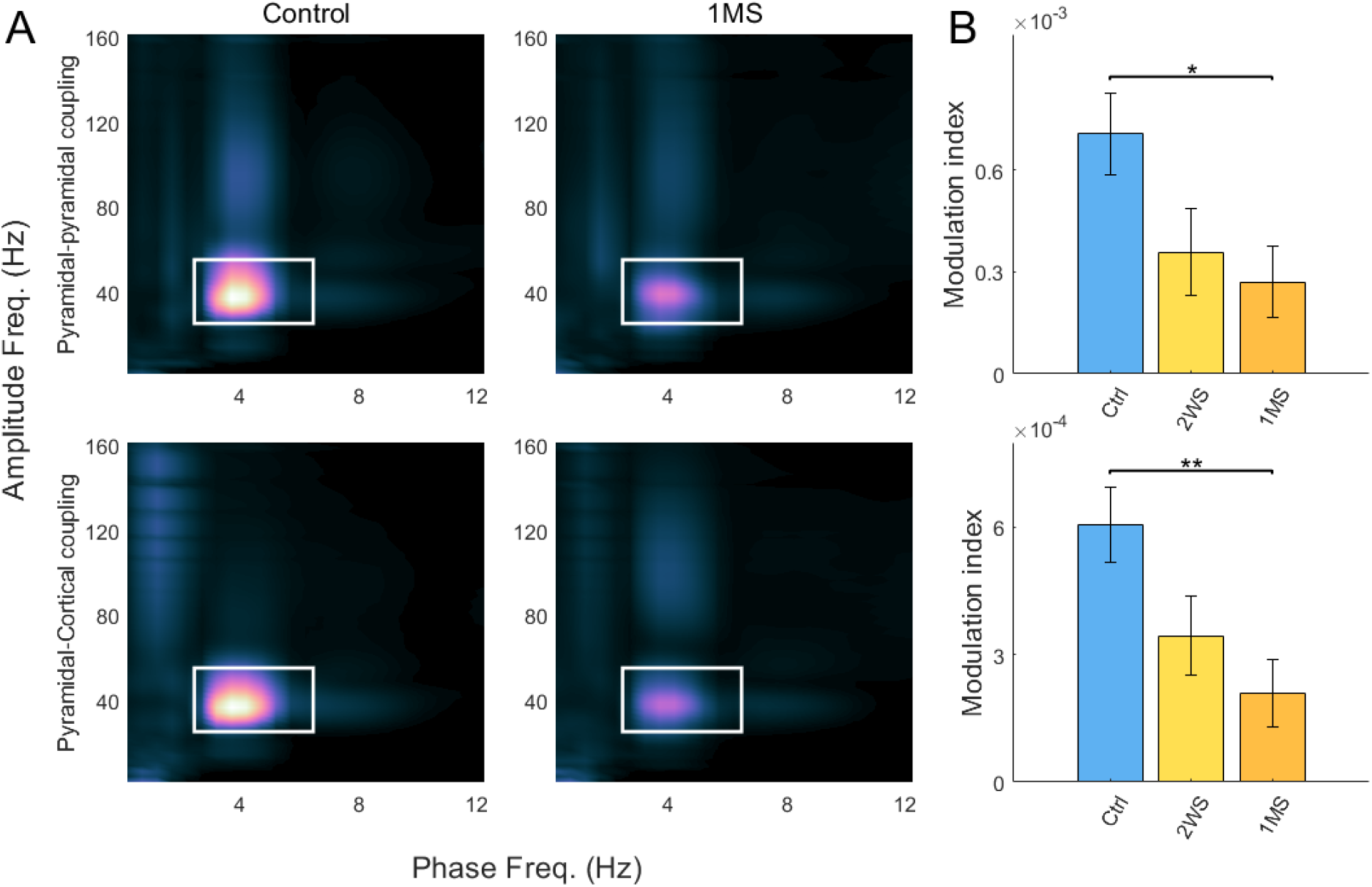
Coupling between theta and gamma. (A) Phase amplitude comodulograms displaying modulation within the pyramidal layer (top), and between cortex and hippocampus (bottom). Theta-gamma coupling demarked by white rectangle. (B) Comparison of average modulation index between theta and gamma. Ipsilesional and contralesional hemispheres are compared separately. *Significant differences (p < 0.05 and p < 0.01, are demarked with * and ** respectively).*

### The effect of Environmental Enrichment on stroke

Following our analysis of stroke progression, we investigated the effect of EE on the hippocampal electrophysiological biomarkers using a two-way ANOVA. EE had two main interactions with the biomarkers affected by stroke. Characteristics of SPW-Rs, which were increased by stroke, were mitigated by EE. However, interestingly, other biomarkers that were disrupted by stroke, such as TD state and PAC, were further disrupted by EE.

To analyze the effect of EE on hippocampal biomarkers, we first looked at characteristics of SPW-Rs. We started with SPW-R power. At 1MS SPW-R power is significantly higher than control in both hemispheres, (Figure 6A) (ANOVA; ipsilesional p = 2.16e-4, contralesional p = 1.06e-24), while there is no significant difference between control, EES, and EEC (ANOVA; p > 0.058). SPW-R power show that EE mitigates the effects following stroke.

**Figure 6.**
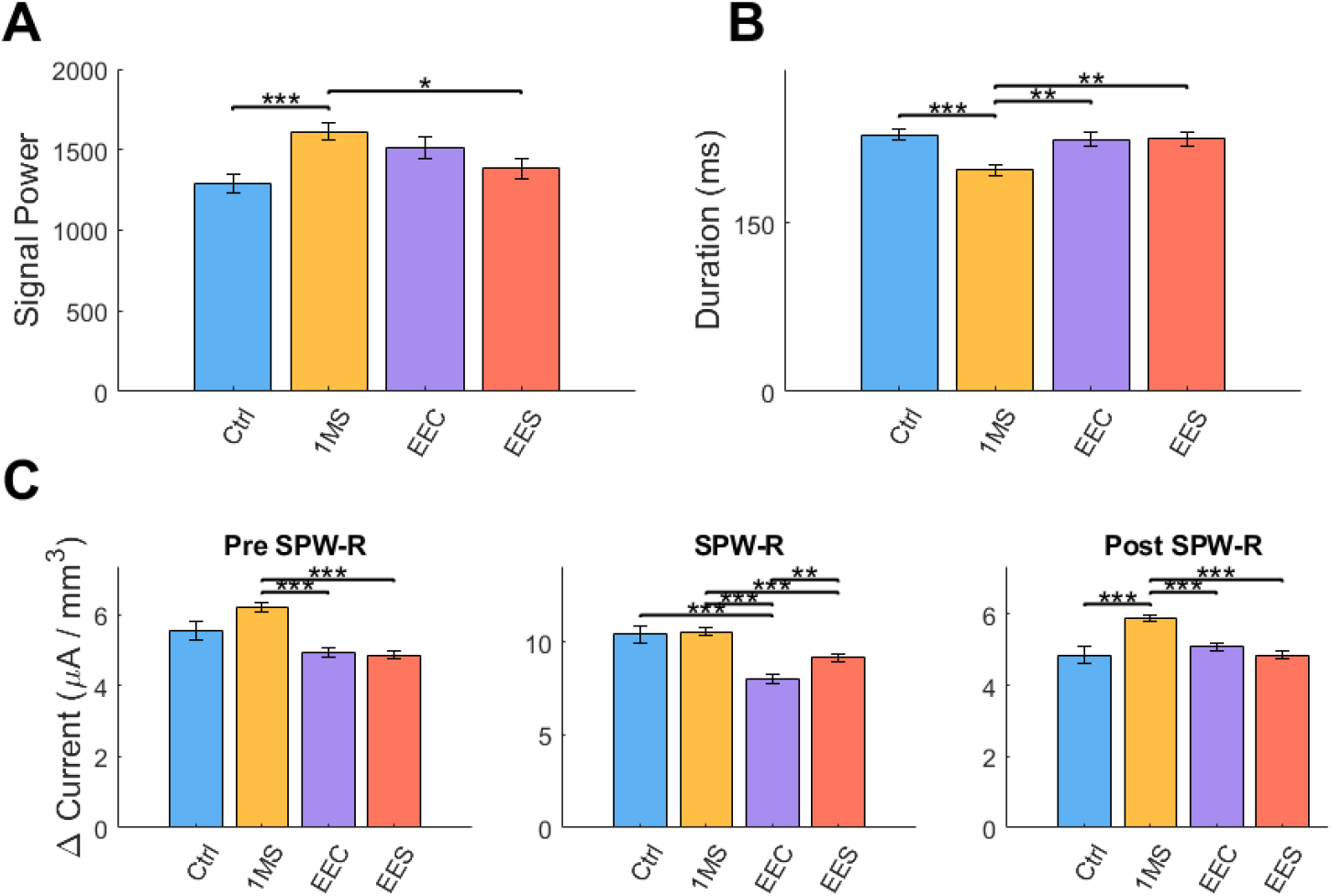
Summary of the effects of EE following stroke on SPW-R characteristics using 2-way ANOVA comparisons on ipsilesional hemisphere. (A) Changes in SPW-R power (B) Changes in SPW-R power. (C) Changes in CSD dipole amplitude surrounding SPW-R. *Significant differences: p < 0.05, p < 0.01 andp < 0.001, are demarked with *, **, or *** respectively.*

We then analyzed the duration of SPW-Rs. At 1MS the duration of SPW-Rs is significantly shorter than control ipsilesionally (ANOVA; p = 5.65e-5), while control, EES, and EEC are not significantly different (ANOVA; p = 1) (Figure 6B). Contralesionally, the duration of 1MS, EEC, and EES are all significantly shorter than control (Supplementary Figure 7B) (ANOVA; 1MS p = 1.98e-25, EEC p = 2.84e-13, EES p = 2.61e-10), though 1MS is also significantly shorter than EES. Like SPW-R power, SPW-R duration results show that EE mitigates the decrease in duration following stroke. These results support our findings in SPW-R power that EE tends to reduce the severity of the effects of stroke.

As for the effects on the CSD surrounding SPW-Rs, we see that stroke generally causes an increase in dipole amplitude, while EE generally causes a decrease in dipole amplitude. Leading up to SPW-R, there is a between-subjects effect in both the ipsilesional and contralesional hemisphere for stroke (ANOVA; ipsilesional p = 3.64e-8, contralesional p = 2.09e-6), (ANOVA; p = 0.032) (Supplementary Table 1). The dipole amplitude ipsilesionally at 1MS is significantly higher than EEC and EES before, during, and after SPW-R (ANOVA; before; EEC p = 4.35e-11, EES p = 3.22e-14, during; EEC p = 6.50e-15, EES p = 1.77e-5, after; EEC p = 4.67e-6, EES p = 4.85e-11), while there is no significant difference between control, EEC, and EES (ANOVA, p > 0.15) (Figure 6C). These changes support our findings that EE mitigates the effects of stroke.

Investigating the effect of EE on TD states revealed a between-subjects effect that both stroke and EE significantly decrease the length of ipsilesional HTD state (ANOVA; stroke p = 4.82e-5, EE p = 0.0041), though contralesionally, only stroke significantly changed HTD state (ANOVA; p = 3.01e-6) (Supplementary Table 1). This was shown in our post-hoc analysis as well, where the length of ipsilesional HTD in 1MS, EEC, and EES were all significantly lower than control (ANOVA; 1MS p = 6.91e-4, EEC p = 0.021, EES p = 3.70e-5) (Figure 7A). Neither stroke nor EE had any significant effect on the ratio of HTD/LTD (Figure 7B, Table 1). These results show that both EE and stroke can disrupt the stability of TD states, while leaving the ratio of HTD/LTD intact.

**Figure 7.**
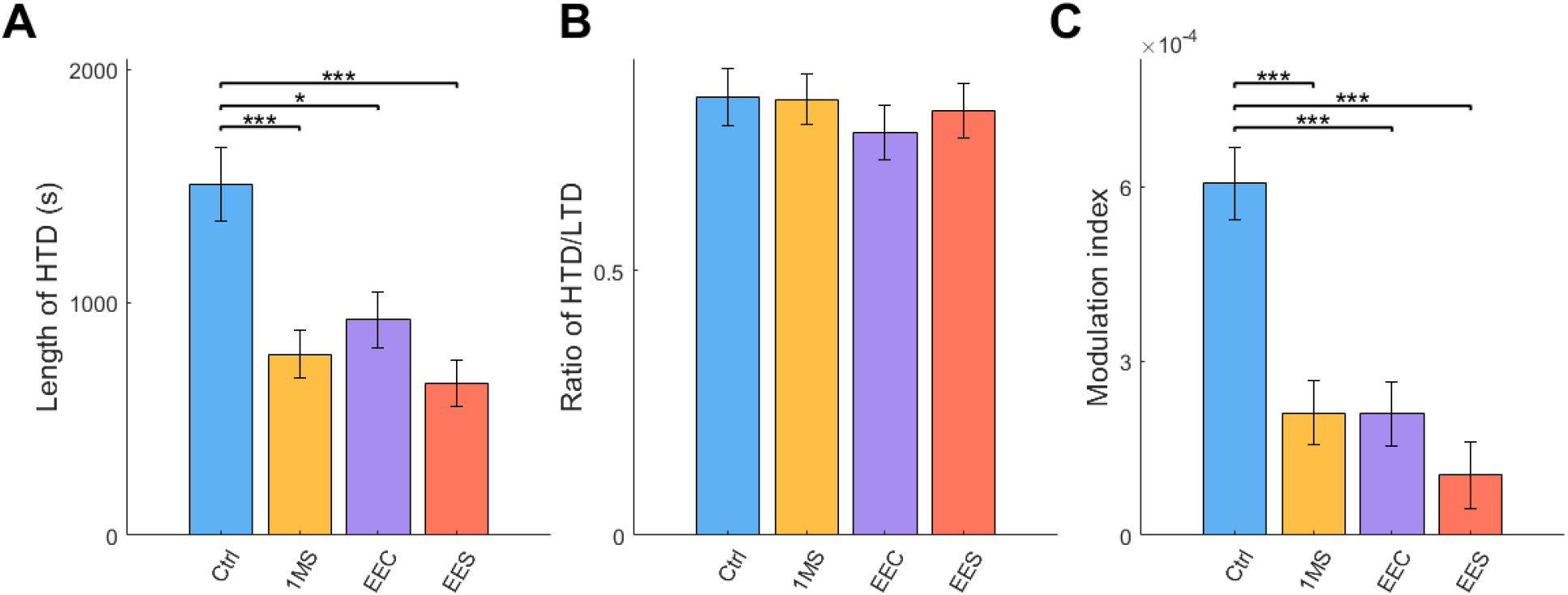
The effects of EE following stroke on HTD state and PAC using 2-way ANOVA comparisons on ipsilesional hemisphere. (A) Change in average HTD state (B) Change in HTD/LTD ratio. (C) Changes in theta-gamma coupling between cortex and pyramidal. *Significant differences: p < 0.05, p < 0.01 andp < 0.001, are demarked with *, **, or *** respectively*.

Contrary to its known benefit on synaptic plasticity and cognition, EE unexpectedly lowered the levels of theta-gamma coupling during HTD. The between-subjects effects show both stroke and EE significantly lower ipsilesional and contralesional theta-gamma coupling. Additionally, stroke and EE showed significant interaction ipsilesionally, meaning that the change in PAC seen in EES compared to control was significantly different than could be expected from the additive effects of stroke and EE combined (ANOVA; ipsilesional; stroke p = 1.21e-4, EE p = 1.14e-4, interaction p = 0.017) (Figure 7C, Table 1). Our post-hoc analysis revealed that coupling in 1MS, EEC, and EES are all significantly lower than control ipsilesionally (ANOVA; 1MS p = 2.10e-4, EEC p = 2.02e-4, EES p = 7.04e-6), while contralesionally coupling in 1MS and EES are significantly lower than control (ANOVA; 1MS p = 0.0022, EES p = 4.88e-4) (Supplementary Figure 8C). These results additionally show a reduction of information flow between cortico-hippocampal networks for both stroke and EE groups.

## Discussion

While impairment of memory after dMCAO is well reported, e.g. poor performance in the Barnes Maze test and hippocampal hypoactivation following spatial exploration (Wang 2011), the electrophysiological substrates of cognitive deficit in the hippocampus have not been established. There are no direct projections between sensorimotor cortex and hippocampus. However, we recently showed that cortical lesion following dMCAO stroke acutely affects the electrophysiology of the hippocampus. We saw counterintuitive effects, such as an increase in aberrant SPW-Rs, an increase in theta-gamma coupling, and a persistent increase in LTD state (He 2019). These results showed that MCA stroke strongly affects distant regions like hippocampus. With these putative biomarkers, we sought to understand the underlying changes to hippocampal electrophysiology that drive the cognitive deficit observed during chronic phase of stroke, and how EE interacts with these effects.

We found that EE mitigates the stroke induced changes to SPW-R characteristics, like SPW-R power, duration, and CSD. This shows for the first time that although the anatomy of hippocampus is not compromised, SPW-Rs, which are well known and thoroughly studied for their role in memory and cognition, are disrupted throughout stroke progression. Crucially, we show that EE stabilizes the characteristics of SPW-Rs throughout stroke progression, revealing that EE impacts biomarkers related to cognition. These results begin to uncover the complex interaction between stroke and EE, providing a testbed to assess different strategies for therapeutics following stroke. Our recordings allowed analysis of other biomarkers as well, though their role in cognition and memory is less clear. For these biomarkers such as TD brain state and PAC, EE seems to compound the disruptions cause by stroke. These biomarkers may be areas of interest to further understand the network imbalances caused by stroke and to investigate future therapeutic interventions.

Current literature describes stroke progression as two opposing phases. The first phase lasts until approximately three days after onset, and is characterized by increased activity and plasticity, as well as excitotoxic cell death. Following this increase in activity, neuronal activity is chronically suppressed (Carmichael 2012). Our results show that SPW-Rs, which encode memories within the hippocampus, can remain upregulated for as long as two weeks before switching phases to a suppressed state, while other biomarkers such as TD state and PAC are chronically disrupted.

The chronic changes to electrophysiology differ drastically from the acute effects of ischemia occurring in the hour after infarct, detailed in (He 2019), such as SPW-Rs the frequency of SPW-Rs increasing in the acute setting, while in the chronic setting SPW-R power increases at 2WS, which may indicate a larger population of recruited neurons firing in each SPW-R (Schlinghoff 2014). PAC also differs between acute and chronic settings, where theta gamma coupling is increased in the acute setting and disrupted in the chronic setting.

Our current results, in conjunction with our previous findings (He 2019), suggest that SPW-Rs respond robustly to stroke. SPW-Rs are well-known for their causative role in memory performance: disruption on SPW-Rs interferes with memory formation (Buzsaki 2015). The marked increase in SPW-R power, duration, and current flow at 2WS may be correlated with the increased cortical plasticity or disinhibition during stroke progression. This suggests some compensatory activity at 2WS. The decrease in SPW-R power, duration, and current flow at 1MS could then represent stabilization of the network. Recent work has shown that long-duration SPW-Rs are correlated with increased memory function (Fernandez-Ruiz et al, 2019), which may imply that shorter SPW-Rs impair memory. Our current results as well as our previous findings that dMCAO impairs cognition and spatial memory (Wang 2011) agrees with this interpretation.

The other biomarkers that we analyzed, such as TD state and PAC were lower at 2WS and further disrupted at 1MS compared to controls. Disrupted TD states have been shown to cause neuroinflammation and have been associated with impairment of learning and memory (Williams and Tortella 2002, Williams 2003, Zhu 2012, Aminov 2017, Ip 2019), which may be a contributing mechanism to post-stroke cognitive impairment. PAC within the hippocampus has been shown as a mechanism for memory processing during sleep, which is correlated with multiple phenomena such as neocortical slow oscillations, and thalamo-cortical sleep spindles (Fell and Axmacher, 2011; Rasch and Born, 2013; Staresina 2015; Bergmann & Born 2018). The breakdown of hippocampal theta rhythms, which are known to coordinate oscillations in many regions, such as entorhinal cortex and prefrontal cortex, may be a contributing factor to cognitive impairment following stroke.

EE has been shown to consistently improve behavioral measures of cognitive recovery following stroke (Hamm 1996; Passineau 2001; Ip 2002; Matsumori 2006; Wang 2011; Wang 2019). We show that EE stabilizes the characteristics of SPW-Rs throughout stroke progression. However, the other biomarkers we analyzed did not reflect this effect. TD state and PAC are further disrupted by EE following stroke. TD states, though measured in the hippocampus, are an indicator for functions of many areas of the brain. Theta-gamma coupling, which we observed within the hippocampus and between cortex and hippocampus, has also been reported between prefrontal cortex and entorhinal cortex. Therefore, the biomarkers which are further disrupted by EE are indicators of a more global effect on the brain compared to SPW-Rs, that occur locally, which may explain why they react differently. These biomarkers may be areas of interest for future therapeutic intervention to improve behavioral outcomes.

This study is limited in that our dataset consists of single time point recordings that were done under urethane anesthesia. While general anesthetics are known to reduce spike activity (Suzuki and Smith 1988), urethane anesthesia has been shown to preserve brain rhythms of interest and generate naturalistic sleep, even inducing spontaneous oscillations (Kramis 1975, Hara and Harris 2002, Pagliardini 2013). Urethane anesthesia is widely used both in hippocampal studies (Klausberger and Somogyi 2008), as well as stroke studies (Rabiller 2015, Moyanova and Dijkhuizen 2014, Srejic 2013). However, a single recording time point prevents assessment of neurophysiology over time and limits our observations to between-group analysis of different animals. An awake behaving recording setup will allow us to gain a more complete understanding of how stroke affects hippocampal electrophysiology across time. Another limitation of unconscious recordings is they do not provide real-time correlates to spatial encoding or recall, or hippocampal activity before and after novel stimuli. An awake-behaving set-up would allow us to record electrophysiology during these events and pair them with the downstream behavioral readouts.

Stroke causes complex changes to many remote regions of the brain beyond the direct infarct. Many neurological disorders are not isolated to a single area, having far reaching effects in remote brain regions. To create effective therapies for these disorders, a deeper understanding of how these remote regions interact is needed. Furthermore, developing a greater understanding of how cognitive therapies like EE affect networks of the brain is essential to develop and translate therapies to clinical settings. Here we have shown that EE impacts SPW-Rs that can lead to cognitive improvement following stroke, however the effects on other biomarkers such as PAC require further study. For example, understanding the changes to PAC following stroke can uncover insights that may translate to many neurological disorders; for example, abnormal PAC has been implicated for many disorders, such Parkinson’s disorder (Devergnas et al 2019), Alzheimer’s Disease (Zhang 2016), and schizophrenia (Barr 2017). This understanding will open the door for more targeted therapies as well. Recently we have shown that PAC can be induced through optogenetic stimulation (Yazdan-Shahmorad 2018), which allows for the potential to recover PAC between regions in disease models.

## Materials & Methods

### A. Animals

We conducted all experiments in accordance with the animal care guidelines issued by the National Institutes of Health and by the San Francisco VA Medical Center Institutional Animal Care and Use Committee. A total of 52 adult male Sprague-Dawley rats approximately 2.5 months of age weighing 250g (Charles River Laboratories, Wilmington, MA) were used and housed in institutional standard cages (2 rats per cage) on a 12-hr light/12-hr dark cycle with ad libitum access to food and water before the experimental procedures. The identity of the test subject was blinded to investigators who performed the stroke surgery and recording.

### B. Experimental Stroke

Stroke was induced unilaterally in rats by the dMCAO method in combination with supplemental proximal artery occlusion of the bilateral common carotid arteries (CCAs) under isoflurane (1.5%)/O2(30%)/N2O(68.5%) anesthesia as described previously (Sun H 2011, He 2019), producing cortical infarct restricted to the somatosensory cortex (Wang 2011). In brief, a 1.5 mm diameter burr hole 1 mm rostral to the anterior junction of the zygoma and temporalis bone was made with a dental drill. The dura mater was carefully pierced with a 30-gauge needle. The main trunk of the left MCA was ligated permanently above the rhinal fissure with a 10-0 suture, and the bilateral CCAs were occluded temporarily for 60 min with 4-0 sutures. The sutures over CCAs were then removed to restore blood flow, and the cervical incision was closed. Core temperature was maintained at 37±0.5 °C with a heating blanket and rectal thermistor servo loop throughout the procedure.

### C. Environmental Enrichment

EE therapy was used to evaluate its potential to affect post-stroke electrophysiology. Immediately following MCAO, we randomly assigned rats into EE or standard housing groups. One week after surgery, we transferred the EE group rats to enriched environment cages (dimensions: 76 × 56 × 77 cm; a 2-story cage equipped with a running wheel for spontaneous exercise, a 3-dimensional labyrinth, bedding, a ladder, a house, chains, a hammock, wooden blocks, and nylon bones; 10 rats per cage) for an additional 3 weeks of residence. Similarly, non-stroke control animals assigned to EE treatment were placed in enriched environmental cages for 3 weeks before recording. We changed the arrangement of movable objects once a week to maintain novelty (Matsumori 2006, Wang 2011). Rats assigned to the standard housing groups remained in the institutional standard cages.

### D. Recording

We performed electrophysiological recordings using two 16-site extracellular silicon probes (NeuroNexus Technologies) under urethane anesthesia for two hours (Sigma, 15 mg/kg i.p.). Following craniotomy, 2 electrodes (A1×16-5mm-100-703) were slowly inserted into each hemisphere after the dura mater was pierced to target the dorsal hippocampus at [AP: −3.3 mm; ML: +/− 2 mm] via a stereotaxic frame (David Kopf Instruments, Tujunga, CA, USA) (Figure 1). Real-time data display and an audio aid were used to facilitate the identification of proper recording location while advancing electrodes until characteristic signals from stratum pyramidal and stratum radiatum were detected and recorded. A 2-hr multi-channel recording from bilateral sensorimotor cortex and dorsal hippocampus was collected from each rat. Data were stored at a sampling rate of 32 kHz after band-pass filtering (0.1-9 kHz) with an input range of ± 3 mV (Digital Lynx SX, Neuralynx, USA). All recordings were down sampled to 1250 Hz (Matlab, MathWorks, USA) prior to analysis.

### E. Tissue preparation and infarct assessment

After recording, rats were perfused transcardially with 4% paraformaldehyde in 0.1M phosphate buffer, pH 7.4. The brains were collected, post fixed overnight in 4% PFA and placed in 30% sucrose solution for 24 h. Brains were cut coronally in 40 μm-thick sections and stored at 4°C. Serial coronal sections were stained using the H&E method. Infarct volume was measured by subtracting the difference between intact tissue in the ipsilesional side from the contralesional side using Stereoinvestigator software (Microbrighfield, VA). We determined both the infarct volume and the ratio of infarct to intact tissue volume (Sun 2013).

### F. Data Analysis

We used local field potentials (LFP) from deep cortical layers and four layers from CA1 field hippocampus (stratum oriens, pyramidal, radiatum and lacunosum-moleculare (SLM)) in our analysis. We isolated brain waves from the LFPs by band-pass filtering the following frequency ranges: delta (0.1-3 Hz), theta (4-7 Hz), alpha (7-13 Hz), beta (13-30 Hz), gamma (30-58 Hz), and high-gamma (62-200 Hz). Out of the 52 rats used in this study, we excluded the data from 8 rats after screening for bad channels. The groups had the following counts: control (n = 8), EEC (n = 10), 2WS (n = 7), 1MS (n = 10), and EES (n = 9). To analyze changes to signal power we normalized data by subtracting the mean and dividing by the standard deviation to account for impedance differences between individual electrodes.

To estimate LTD and HTD brain states we calculated the ratio of spontaneous signal power between theta band and delta band from the pyramidal layer. The threshold defining LTD and HTD states was defined manually for each animal by visual assessment (Bodizs 2001, Buzsaki 2002, Karlsson and Frank, 2009).

SPW-Rs were identified when a pyramidal ripple and radiatum sharp wave co-occurred (Karlsson and Frank, 2009, Buzsaki 2015). To detect pyramidal ripples, the LFP of the pyramidal layer was bandpass filtered (150-250 Hz), then squared and Z-scored. When the signal exceeded 6 standard deviations for a period longer than 20 ms, an event was registered. When the signal subsequently dropped below 1 SD, the event was considered ended. To identify radiatum sharp waves, a similar process was used, however the bandpass filter was from 8 to 40 Hz, and the standard deviation threshold was 3.

We performed laminar current-source density (CSD) analysis (Kenan-Vaknin and Teyler, 1994) along each electrode, temporally aligning the LFP to the onset of a SPW-R, and spatially centering each recording on the pyramidal layer. The CSD consists of a one-dimensional surface Laplacian along the length of the electrode to approximate the relative sources and sinks through the cortex and hippocampus from one second before ripple onset to one second after. Dipole amplitude was calculated by finding the maximum and minimum current along the probe from the specified time window and taking the difference.

We analyzed PAC within the hippocampus and between the layers of the hippocampus and the cortex as a metric of functional connectivity and communication. PAC was calculated as described in (Tort 2010). Briefly, the LFP was bandpass-filtered between (0.1-200 Hz). The instantaneous phase and amplitude were extracted using the Hilbert transform. A composite phase-amplitude time series then determined the amplitude distribution across phase. The modulation index (MI) is then calculated from the divergence of the amplitude distribution from a uniform distribution (Tort 2010).

### G. Statistical analysis

We expressed data as mean ±standard error. We performed one-way ANOVA to assess changes in stroke progression, and two-way ANOVA to assess changes between the effect of stroke and the effect of EE. We used post-hoc Bonferroni’s to control for multiple comparisons. We performed paired t-tests to assess changes between hemispheres. For non-normal distributions, we performed Kruskal-Wallis with post-hoc Bonferroni’s to control for multiple comparisons. We considered *p* values less than 0.05 as significant.

## Conflict of interest statement

The authors declare no competing financial interests.

## Acknowledgments

This project was supported by the Eunice Kennedy Shriver National Institute of Child Health & Human Development of the National Institutes of Health under Award Number K12HD073945, the Center for Neurotechnology (CNT, a National Science Foundation Engineering Research Center under Grant EEC-1028725), NIH R01 NS102886, VA Merit Award I01BX003335 and VA Research Career Scientist award IK6BX004600. We thank Loren Frank, Kenny Kay and Karam Khateeb for advice on data analysis.

## Notes

### Competing Interest Statement

The authors have declared no competing interest.

## References

Aminov, A., Rogers, J. M., Johnstone, S. J., Middleton, S., & Wilson, P. H. (2017). Acute single channel EEG predictors of cognitive function after stroke. PLoS ONE, 12(10), 1–15. https://doi.org/10.1371/journal.pone.0185841

Bandarabadi, M., Boyce, R., Herrera, C. G., Bassetti, C. L., Williams, S., Schindler, K., & Adamantidis, A. (2019). Dynamic modulation of theta-gamma coupling during rapid eye movement sleep. Sleep, 42(12), 1–11. https://doi.org/10.1093/sleep/zsz182

Barker-Collo, S., Feigin, V., Lawes, C., Parag, V., & Senior, H. (2010). Attention deficits after incident stroke in the acute period: Frequency across types of attention and relationships to patient characteristics and functional outcomes. Topics in Stroke Rehabilitation, 17(6), 463–476. https://doi.org/10.1310/tsr1706-463

Barr, M. S., Rajji, T. K., Zomorrodi, R., Radhu, N., George, T. P., Blumberger, D. M., & Daskalakis, Z. J. (2017). Impaired theta-gamma coupling during working memory performance in schizophrenia. Schizophrenia Research, 189, 104–110. https://doi.org/10.1016/j.schres.2017.01.044

Barth, A. M. I., & Mody, I. (2011). Changes in hippocampal neuronal activity during and after unilateral selective hippocampal ischemia in vivo. Journal of Neuroscience, 31(3), 851–860. https://doi.org/10.1523/JNEUROSCI.5080-10.2011

Battaglia, F. P., Benchenane, K., Sirota, A., Pennartz, C. M. A., & Wiener, S. I. (2011). The hippocampus: Hub of brain network communication for memory. Trends in Cognitive Sciences, 15(7), 310–318. https://doi.org/10.1016/j.tics.2011.05.008

Bederson Joshua, B., Pitts Lawrence, H., Tsuji Miles, C., Nishimura Merry, L., Davis Richard, L., & Bartkowski Henry, L. (1986). Rat Middle Cerebral Artery Occlusion: Evaluation of the Model and Development of a Neurologic Examination. Stroke, 17(3), 472–476. https://doi.org/10.1161/01.STR.17.3.472

Bergmann, T. O., & Born, J. (2018). Phase-Amplitude Coupling: A General Mechanism for Memory Processing and Synaptic Plasticity? Neuron, 97(1), 10–13. https://doi.org/10.1016/j.neuron.2017.12.023

Bódizs, R., Kántor, S., Szabó, G., Szûcs, A., Erõss, L., & Halász, P. (2001). Rhythmic hippocampal slow oscillation characterizes REM sleep in humans. Hippocampus, 11(6), 747–753. https://doi.org/10.1002/hipo.1090

Buzsaki, G. (2015). Hippocampal Sharp Wave-Ripple: A Cognitive Biomarkerfor Episodic Memory and Planning. Hippocampus, 25(10). https://doi.org/https://doi.org/10.1002/hipo.22488

Buzsáki, G. (2002). Theta oscillations in the hippocampus. Neuron, 33(3), 325–340. https://doi.org/10.1016/S0896-6273(02)00586-X

Carmichael, S. T. (2012). Brain excitability in stroke: The yin and yang of stroke progression. Archives of Neurology, 69(2), 161–167. https://doi.org/10.1001/archneurol.2011.1175

Carr, M. F., Jadhav, S. P., & Frank, L. M. (2011). Hippocampal replay in the awake state: A potential substrate for memory consolidation and retrieval. Nature Neuroscience, 14(2), 147–153. https://doi.org/10.1038/nn.2732

Cooper, R. M., and Zubek, J. P. (1958). Effects of enriched and restricted early environments on the learning ability of bright and dull rats. Can. J. Psychol. 12, 159–164.

Colgin, L. L. (2015). Theta-gamma coupling in the entorhinal-hippocampal system. Current Opinion in Neurobiology, 31, 45–50. https://doi.org/10.1016/j.conb.2014.08.001

Cumming, T. B., Plummer-D’Amato, P., Linden, T., & Bernhardt, J. (2009). Hemispatial Neglect and Rehabilitation in Acute Stroke. Archives of Physical Medicine and Rehabilitation, 90(11), 1931–1936. https://doi.org/10.1016/j.apmr.2009.04.022

Devergnas, A., Caiola, M., Pittard, D., & Wichmann, T. (2019). Cortical Phase-Amplitude Coupling in a Progressive Model of Parkinsonism in Nonhuman Primates. Cerebral Cortex, 29(1), 167–177. https://doi.org/10.1093/cercor/bhx314

Diamond, M., Law, F., Rhodes, H., Lindner, B., Rosenzweig, M., Krech, D., & Bennett, E. (1966). Increases in cortical depth and glia numbers in rats subjected to enriched environment. Journal of Comparative Neurology, 128(1), 117–125.

Fan, Y., Liu, Z., Weinstein, P. R., Fike, J. R., & Liu, J. (2007). Environmental enrichment enhances neurogenesis and improves functional outcome after cranial irradiation. European Journal of Neuroscience, 25(1), 38–46. https://doi.org/10.1111/j.1460-9568.2006.05269.x

Fernández-Ruiz, A., Oliva, A., de Oliveira, E. F., Rocha-Almeida, F., Tingley, D., & Buzsáki, G. (2019). Long-duration hippocampal sharp wave ripples improve memory. Science, 364(6445), 1082–1086. https://doi.org/10.1126/science.aax0758

Hamm, R. J, Temple, M. D, O’Dell, D. M, Pike, B. R, & Lyeth, B. G. (1996). Exposure to environmental complexity promotes recovery of cognitive function after traumatic brain injury. Journal of Neurotrauma, 13(1), 41–47.

Hanslmayr, S., Staresina, B. P., & Bowman, H. (2016). Oscillations and Episodic Memory: Addressing the Synchronization/Desynchronization Conundrum. Trends in Neurosciences, 39(1), 16–25. https://doi.org/10.1016/j.tins.2015.11.004

Hara, K., & Harris, R. A. (2002). The anesthetic mechanism of urethane: The effects on neurotransmitter-gated ion channels. Anesthesia and Analgesia, 94(2), 313–318. https://doi.org/10.1213/00000539-200202000-00015

Hazime, M., Alasoadura, M., Lamtahri, R., Quilichini, P., Leprince, J., Vaudry, D., & Chuquet, J. (2020). Prolonged deficit of gamma oscillations in the peri-infarct cortex of mice after stroke, https://doi.org/10.1101/2020.03.05.978593

He, J. W., Rabiller, G., Nishijima, Y., Akamatsu, Y., Khateeb, K., Yazdan-Shahmorad, A., & Liu, J. (2019). Experimental cortical stroke induces aberrant increase of sharp-wave-associated ripples in the hippocampus and disrupts cortico-hippocampal communication. Journal of Cerebral Blood Flow and Metabolism. https://doi.org/10.1177/0271678X19877889

Heusser, A. C., Poeppel, D., Ezzyat, Y., & Davachi, L. (2016). Episodic sequence memory is supported by a theta-gamma phase code. Nature Neuroscience, 19(10), 1374–1380. https://doi.org/10.1038/nn.4374

Iizuka, H., Sakatani, K., & Young, W. (1989). Selective cortical neuronal damage after middle cerebral artery occlusion in rats. Stroke, 20(11), 1516–1523. https://doi.org/10.1161/01.STR.20.11.1516

Ip, Z., Rabiller, G., He, J. W., Yao, Z., Akamatsu, Y., Nishijima, Y., Liu, J., & Yazdan-Shahmorad, A. (2019). Cortical stroke affects activity and stability of theta/delta states in remote hippocampal regions *. 5225–5228. https://doi.org/10.1109/embc.2019.8857679

Ip, E Y, Giza, C C, Griesbach, G S, & Hovda, D A. (2002). Effects of enriched environment and fluid percussion injury on dendritic arborization within the cerebral cortex of the developing rat. Journal of Neurotrauma, 19(5), 573–585.

Jadhav, S. P., Kemere, C., German, P. W., & Frank, L. M. (2012). Awake Hippocampal SharpWave Ripples Support Spatial Memory. Science, 336(6087), 1454–1458. https://doi.org/10.1126/science.1217230.Awake

Karlsson, M. P., & Frank, L. M. (2009). Awake_replay_of_remote_experie.PDF. Nature Neuroscience, 12(7), 913–918.

Kenan-Vaknin, G., & Teyler, T. J. (1994). Laminar pattern of synaptic activity in rat primary visual cortex: comparison of in vivo and in vitro studies employing the current source density analysis. Brain Research, 635(1-2), 37–48. https://doi.org/10.1016/0006-8993(94)91421-4

Kenneth, K., Marielena, S., Chung, J. E., Karlsson, M. P., Larkin, M. C., & Frank, L. M. (2016). A_hippocampal_network_for_spat.PDF. Nature, 531(7593). https://doi.org/http://dx.doi.org.offcampus.lib.washington.edu/10.1038/nature17144

Khedr, E. M., Hamed, S. A., El-Shereef, H. K., Shawky, O. A., Mohamed, K. A., Awad, E. M., Ahmed, M. A., Shehata, G. A., & Eltahtawy, M. A. (2009). Cognitive impairment after cerebrovascular stroke: Relationship to vascular risk factors. Neuropsychiatric Disease and Treatment, 5(1), 103–116. https://doi.org/10.2147/ndt.s4184

Klausberger, T., & Somogyi, P. (2008). Neuronal diversity and temporal dynamics: The unity of hippocampal circuit operations. Science, 321(5885), 53–57. https://doi.org/10.1126/science.1149381

Komitova, M., Mattsson, B., Johansson, B. B., & Eriksson, P. S. (2005). Enriched environment increases neural stem/progenitor cell proliferation and neurogenesis in the subventricular zone of stroke-lesioned adult rats. Stroke, 36(6), 1278–1282. https://doi.org/10.1161/01.STR.0000166197.94147.59

Kramis, R., Vanderwolf, C. H., & Bland, B. H. (1975). Two types of hippocampal rhythmical slow activity in both the rabbit and the rat: Relations to behavior and effects of atropine, diethyl ether, urethane, and pentobarbital. Experimental Neurology, 49(1), 58–85. https://doi.org/10.1016/0014-4886(75)90195-8

Lisman, J., & Buzsáki, G. (2008). A neural coding scheme formed by the combined function of gamma and theta oscillations. Schizophrenia Bu lle tin, 34(5), 974–980. https://doi.org/10.1093/schbul/sbn060

Liu, F., & McCullough, L. D. (2011). Middle cerebral artery occlusion model in rodents: Methods and potential pitfalls. Journal of Biomedicine and Biotechnology, 2011. https://doi.org/10.1155/2011/464701

Manosevitz, M. (1970). Early environmental enrichment and mouse behavior. J. Comp. Physiol. Psychol. 71, 459–466. doi: 10.1037/h0029141

Matsumori, Y., Hong, S. M., Fan, Y., Kayama, T., Hsu, C. Y., Weinstein, P. R., & Liu, J. (2006). Enriched environment and spatial learning enhance hippocampal neurogenesis and salvages ischemic penumbra after focal cerebral ischemia. Neurobiology of Disease, 22(1), 187–198. https://doi.org/10.1016/j.nbd.2005.10.015

McNaughton, N., Ruan, M., & Woodnorth, M.-A. (2006). Restoring Theta-like Rhythmicity in Rats Restores Inital Learning in Morris Water Maze. Hippocampus, 16, 1102–1110. https://doi.org/10.1002/hipo

Moyanova, S. G., & Dijkhuizen, R. M. (2014). Present status and future challenges of electroencephalography- and magnetic resonance imaging-based monitoring in preclinical models of focal cerebral ischemia. Brain Research Bulletin, 102, 22–36. https://doi.org/10.1016/j.brainresbull.2014.01.003

Nilsson, M., Perfilieva, E., Johansson, U., Orwar, O., & Eriksson, P. S. (1999). Enriched environment increases neurogenesis in the adult rat dentate gyrus and improves spatial memory. Journal of Neurobiology, 39(4), 569–578. https://doi.org/10.1002/(SICI)1097-4695(19990615)39:4<569::AID-NEU10>3.0.CO;2-F

Ognjanovski, N., Maruyama, D., Lashner, N., Zochowski, M., & Aton, S. J. (2014). CA1 hippocampal network activity changes during sleep-dependent memory consolidation. Frontiers in Systems Neuroscience, 8(1 APR), 1–11. https://doi.org/10.3389/fnsys.2014.00061

Okada, Mitsuko, Tamura, Akira, Urae, Akinori, Nakagomi, Tadayoshi, Kirino, Takaaki, Mine, Kazunori, & Fujiwara, Michihiro. (1995). Long-Term Spatial Cognitive Impairment Following Middle Cerebral Artery Occlusion in Rats. A Behavioral Study. Journal of Cerebral Blood Flow and Metabolism, 15(3), 505–512.

Oliveira, J. L. de, Crispin, P. di T. B., Duarte, E. C. W., Marloch, G. D., Gargioni, R., Trentin, A. G., & Alvarez-Silva, M. (2014). Histopathology of motor cortex in an experimental focal ischemic stroke in mouse model. Journal of Chemical Neuroanatomy, 57-58, 1–9. https://doi.org/10.1016/j.jchemneu.2014.03.002

Pagliardini, S., Gosgnach, S., & Dickson, C. T. (2013). Spontaneous Sleep-Like Brain State Alternations and Breathing Characteristics in Urethane Anesthetized Mice. PLoS ONE, 8(7), 1–11. https://doi.org/10.1371/journal.pone.0070411

Passineau, M. J., Green, E. J., & Dietrich, W. D. (2001). Therapeutic effects of environmental enrichment on cognitive function and tissue integrity following severe traumatic brain injury in rats. Experimental Neurology, 168(2), 373–384. https://doi.org/10.1006/exnr.2000.7623

Rabiller, G., He, J. W., Nishijima, Y., Wong, A., & Liu, J. (2015). Perturbation of brain oscillations after ischemic stroke: A potential biomarker for post-stroke function and therapy. International Journal of Molecular Sciences, 16(10), 25605–25640. https://doi.org/10.3390/ijms161025605

Rasch, B., & Born, J. (2013). About sleep’s role in memory. Physiological Reviews, 93(2), 681–766. https://doi.org/10.1152/physrev.00032.2012

Schlingloff, D., Káli, S., Freund, T. F., Hájos, N., & Gulyás, A. I. (2014). Mechanisms of sharp wave initiation and ripple generation. Journal of Neuroscience, 34(34), 11385–11398. https://doi.org/10.1523/JNEUROSCI.0867-14.2014

Schmitt, O., Badurek, S., Liu, W., Wang, Y., Rabiller, G., Kanoke, A., Eipert, P., & Liu, J. (2017). Prediction of regional functional impairment following experimental stroke via connectome analysis. Scientific Reports, 7(December 2016), 1–18. https://doi.org/10.1038/srep46316

Shirvalkar, P. R., Rapp, P. R., & Shapiro, M. L. (2010). Bidirectional changes to hippocampal theta-gamma comodulation predict memory for recent spatial episodes. Proceedings of the National Academy of Sciences of the United States of America, 107(15), 7054–7059. https://doi.org/10.1073/pnas.0911184107

Silasi, G., & Murphy, T. H. (2014). Stroke and the Connectome: How Connectivity Guides Therapeutic Intervention. Neuron, 84(2), 511. https://doi.org/10.1016/j.neuron.2014.10.020

Srejic, L. R., Valiante, T. A., Aarts, M. M., & Hutchison, W. D. (2013). High-frequency cortical activity associated with postischemic epileptiform discharges in an in vivo rat focal stroke model: Laboratory investigation. Journal of Neurosurgery, 118(5), 1098–1106. https://doi.org/10.3171/2013.1.JNS121059

Staresina, B. P., Bergmann, T. O., Bonnefond, M., Van Der Meij, R., Jensen, O., Deuker, L., Elger, C. E., Axmacher, N., & Fell, J. (2015). Hierarchical nesting of slow oscillations, spindles and ripples in the human hippocampus during sleep. Nature Neuroscience, 18(11), 1679–1686. https://doi.org/10.1038/nn.4119

Sun, C., Sun, H., Wu, S., Lee, C. C., Akamatsu, Y., Wang, R. K., Kernie, S. G., & Liu, J. (2013). Conditional Ablation of Neuroprogenitor Cells in Adult Mice Impedes Recovery of Poststroke Cognitive Function and Reduces Synaptic Connectivity in the Perforant Pathway. Journal of Neuroscience, 33(44), 17314–17325. https://doi.org/10.1523/JNEUROSCI.2129-13.2013

Sun H, Le T, Chang TT, Habib A, Wu S, Shen F, Young WL, Su H, Liu J (2011) AAV-mediated netrin-1 overexpression increases periinfarct blood vessel density and improves motor function recovery after experimental stroke. Neurobiology of disease 44:73–83.

Suzuki, S. S., & Smith, G. K. (1988). Spontaneous EEG spikes in the normal hippocampus. V. Effects of ether, urethane, pentobarbital, atropine, diazepam and bicuculline. Electroencephalography and Clinical Neurophysiology, 70(1), 84–95. https://doi.org/10.1016/0013-4694(88)90198-8

Tamura, M., Spellman, T. J., Rosen, A. M., Gogos, J. A., & Gordon, J. A. (2017). Hippocampal-prefrontal theta-gamma coupling during performance of a spatial working memory task. Nature Communications, 8(1). https://doi.org/10.1038/s41467-017-02108-9

Tort, A. B. L., Komorowski, R. W., Manns, J. R., Kopell, N. J., & Eichenbaum, H. (2009). Theta-gamma coupling increases during the learning of item-context associations. Proceedings of the National Academy of Sciences of the United States of America, 106(49), 20942–20947. https://doi.org/10.1073/pnas.0911331106

Tort, A. B. L., Komorowski, R., Eichenbaum, H., & Kopell, N. (2010). Measuring phaseamplitude coupling between neuronal oscillations of different frequencies. Journal of Neurophysiology, 104(2), 1195–1210. https://doi.org/10.1152/jn.00106.2010

Wang, C. J., Wu, Y., Zhang, Q., Yu, K. W., & Wang, Y. Y. (2019). An enriched environment promotes synaptic plasticity and cognitive recovery after permanent middle cerebral artery occlusion in mice. Neural Regeneration Research, 14(3), 462–469. https://doi.org/10.4103/1673-5374.245470

Wang, Y., Bontempi, B., Leinekugel, X., Lesburguères, E., Liu, W., Weinstein, P. R., Zhao, J., Abrams, G. M., & Liu, J. (2011). Environmental Enrichment Preserves Cortical Inputs to the Parahippocampal Areas and Reduces Post Stroke Diaschisis. American Journal of Neuroprotection and Neuroregeneration, 3(1), 66–76. https://doi.org/10.1166/ajnn.2011.1027

Williams, A. J., Lu, X. C. M., Hartings, J. A., & Tortella, F. C. (2003). Neuroprotection assessment by topographic electroencephalographic analysis: Effects of a sodium channel blocker to reduce polymorphic delta activity following ischaemic brain injury in rats. Fundamental and Clinical Pharmacology, 17(5), 581–593. https://doi.org/10.1046/j.1472-8206.2003.00183.x

Williams, A. J., & Tortella, F. C. (2002). Neuroprotective effects of the sodium channel blocker RS100642 and attenuation of ischemia-induced brain seizures in the rat. Brain Research, 932(1-2), 45–55. https://doi.org/10.1016/S0006-8993(02)02275-8

Yazdan-Shahmorad, A., Silversmith, D. B., & Sabes, P. N. (2018). Novel techniques for large-scale manipulations of cortical networks in non-human primates. Conference Proceedings:… Annual International Conference of the IEEE Engineering in Medicine and Biology Society. IEEE Engineering in Medicine and Biology Society. Annual Conference, 2018, 5479–5482. https://doi.org/10.1109/EMBC.2018.8513668

Zhang, X., Zhong, W., Brankačk, J., Weyer, S. W., Müller, U. C., Tort, A. B. L., & Draguhn, A. (2016). Impaired theta-gamma coupling in APP-deficient mice. Scientific Reports, 6, 1–10. https://doi.org/10.1038/srep21948

Zhang S, Tong R, Zhang H, Hu X, Zheng X (2006) A pilot studies in dynamic profile of multi parameters of EEG in a rat model of transient middle cerebral artery occlusion. Conference proceedings: Annual International Conference of the IEEE Engineering in Medicine and Biology Society IEEE Engineering in Medicine and Biology Society Annual Conference 1:1181–1184.

Zhu, B., Dong, Y., Xu, Z., Gompf, H. S., Ward, S. A. P., Xue, Z., Miao, C., Zhang, Y., Chamberlin, N. L., & Xie, Z. (2012). Sleep disturbance induces neuroinflammation and impairment of learning and memory. Neurobiology of Disease, 48(3), 348–355. https://doi.org/10.1016/j.nbd.2012.06.022

