## Supplementary material for "Environmental enrichment mitigates stroke-induced change in sharp-wave associated ripple characteristics": Full Supplement (from another version of this preprint: version 1, 2021-02-14)

### Supplementary Figures:

| Biomarker | Ipsilesional p-values | | | Contralesional p-values | | |
|---|---|---|---|---|---|---|
| **Signal Power** | Stroke | EE | Interaction | Stroke | EE | Interaction |
| Cortical δ | **0.0033** | 0.75 | 0.26 | 0.37 | 0.47 | 0.46 |
| Cortical θ | 0.078 | 0.57 | 0.075 | 0.51 | 0.47 | 0.31 |
| Cortical γ | **0.0092** | 0.16 | 0.24 | 0.33 | 0.17 | 0.65 |
| Cortical High γ | **0.0068** | 0.12 | 0.15 | 0.18 | 0.22 | 0.44 |
| Pyramidal δ | **7.60E-04** | 0.43 | 0.085 | 0.2 | 0.7 | 0.2 |
| Pyramidal θ | 0.22 | 0.56 | 0.073 | 0.63 | 0.65 | 0.12 |
| Pyramidal γ | **0.005** | **0.029** | 0.36 | 0.32 | 0.064 | 0.58 |
| Pyramidal High γ | **0.0022** | 0.095 | 0.25 | 0.26 | 0.12 | 0.41 |
| Oriens δ | **0.0072** | 0.067 | 0.25 | 0.69 | 0.39 | 0.34 |
| Oriens θ | 0.67 | 0.78 | 0.18 | 0.11 | 0.28 | 0.28 |
| Oriens γ | 0.27 | **2.40E-04** | 0.3 | 0.67 | **0.0056** | 0.54 |
| Oriens High γ | 0.076 | **0.0066** | 0.16 | 0.97 | 0.051 | 0.31 |
| SLM δ | 0.076 | 0.75 | 0.14 | 0.2 | 0.77 | 0.08 |
| SLM θ | 0.92 | 0.89 | 0.28 | 0.89 | 0.69 | 0.4 |
| SLM γ | 0.15 | 0.12 | 0.33 | 0.2 | 0.1 | **0.037** |
| SLM High γ | 0.07 | 0.44 | 0.5 | 0.29 | 0.37 | **0.034** |
| **TD State** | | | | | | |
| HTD Length | **4.83E-05** | **0.0041** | 0.061 | **3.01E-06** | 0.37 | **0.038** |
| HTD/LTD Ratio | 0.72 | 0.39 | 0.66 | 0.87 | 0.91 | 0.31 |
| **PAC** | | | | | | |
| δ-γ coupling | **1.15E-04** | **1.21E-04** | **0.017** | **0.032** | **1.17E-04** | 0.19 |
| **SPW-R** | | | | | | |
| Power | 0.0992 | 0.9716 | **1.07E-04** | **2.88E-12** | **8.73E-10** | **3.88E-14** |
| Duration | **0.0053** | **0.033** | **0.0044** | **4.86E-10** | **3.00E-03** | **1.32E-17** |
| **CSD** | | | | | | |
| Pre SPW-R | **3.60E-08** | 0.097 | **0.032** | **2.10E-06** | 0.34 | 0.96 |
| SPW-R | **3.86E-10** | **0.032** | 0.082 | **1.12E-07** | 0.2 | **0.0057** |
| Post SPW-R | **0.0092** | **0.01** | **4.43E-05** | 0.23 | 0.76 | 0.78 |

Supplementary Table 1. Summary of two-way ANOVA tests of between-subject effects for all biomarkers. Values less than 0.05 are demarked in bold.

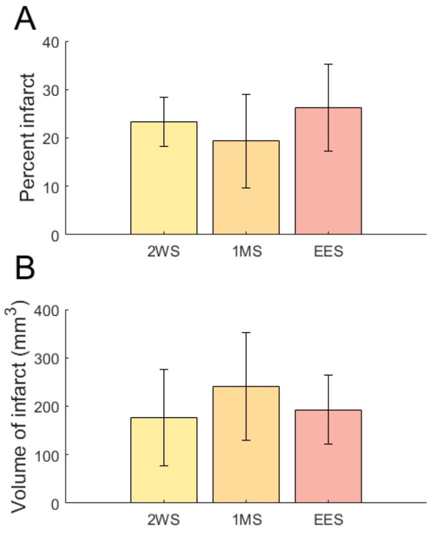


Supplementary Figure 1. Comparison of infarct. (A) Percent of ipsilesional intact tissue to contralesional intact tissue. (B) Differences between ipsilesional intact tissue to contralesional hemisphere.

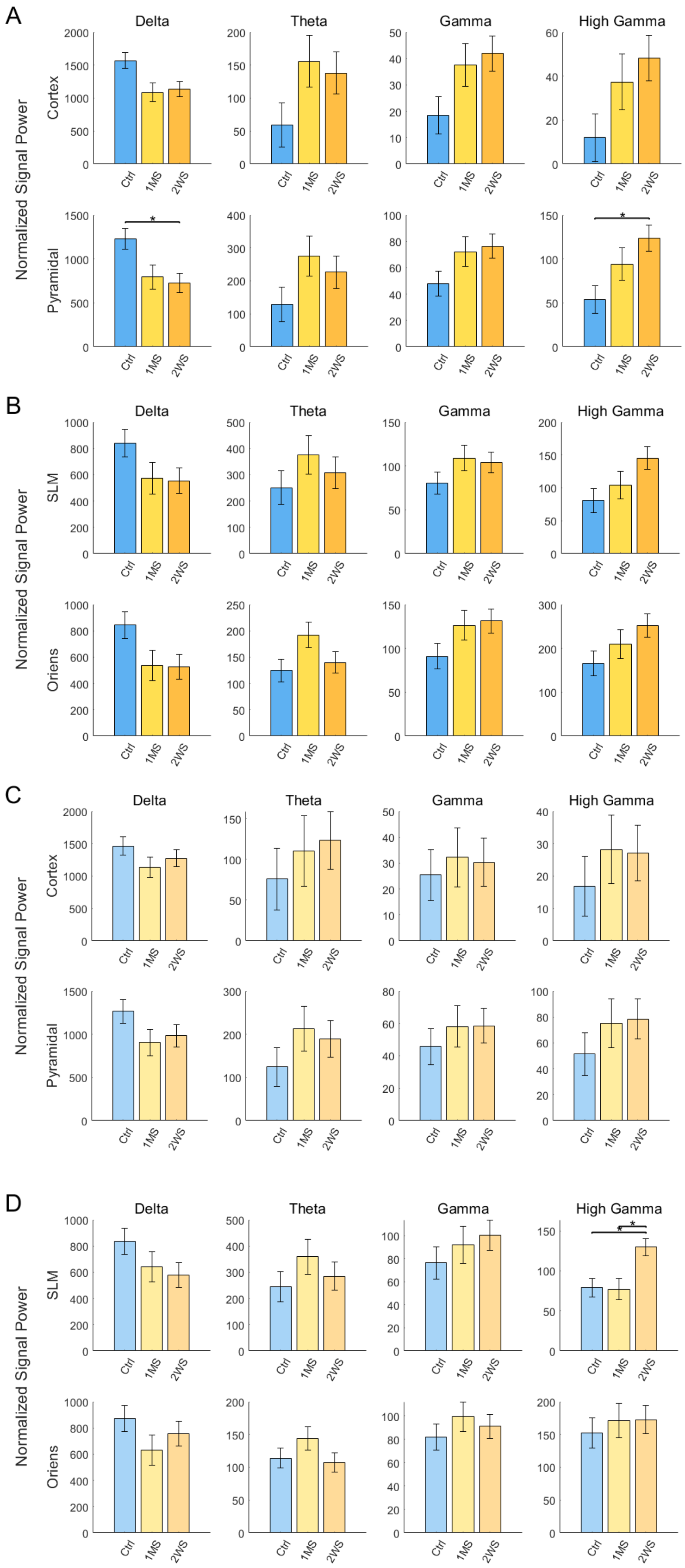
A
B
C
D
Delta
Theta
Gamma
High Gamma
Normalized Signal Power
Cortex
Pyramidal
SLM
Oriens
Ctrl
1MS
2WS
*

Supplementary Figure 2. Comparison of normalized delta, theta, gamma, and high-gamma signal power. (A) Ipsilesional cortex and pyramidal. (B) Ipsilesional SLM and Oriens. (C) Contralesional cortex and pyramidal (D) Contralesional SLM and Oriens. *Significant differences (p < 0.05, are demarked with *).*

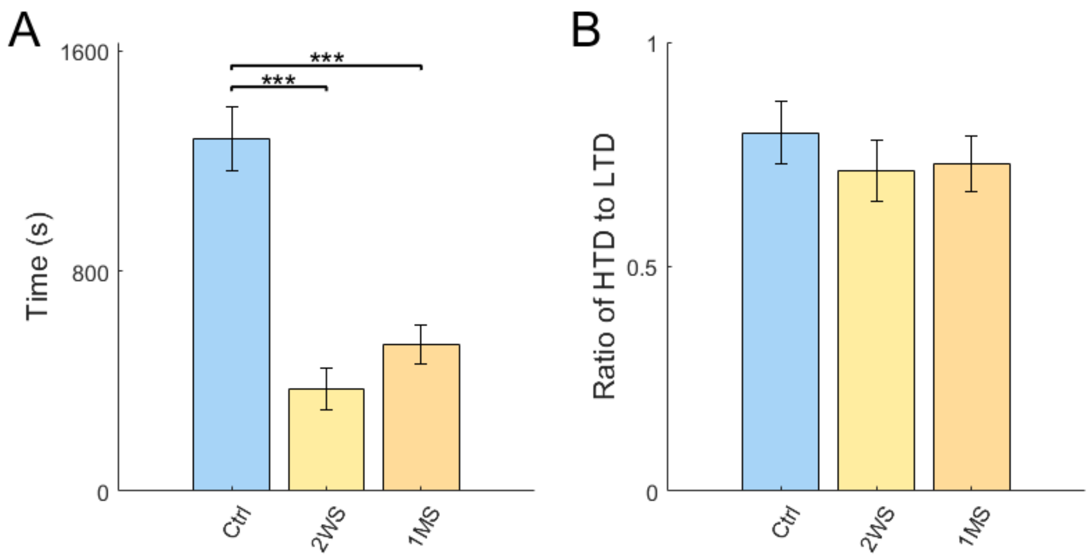


Supplementary Figure 3. (A) Comparison of the average duration of each contralesional HTD state. (B) Comparison of the contralesional proportion of HTD to LTD. *Significant differences (p < 0.05, p < 0.01, p < 0.001, are demarked with *,**,or ***, respectively).*

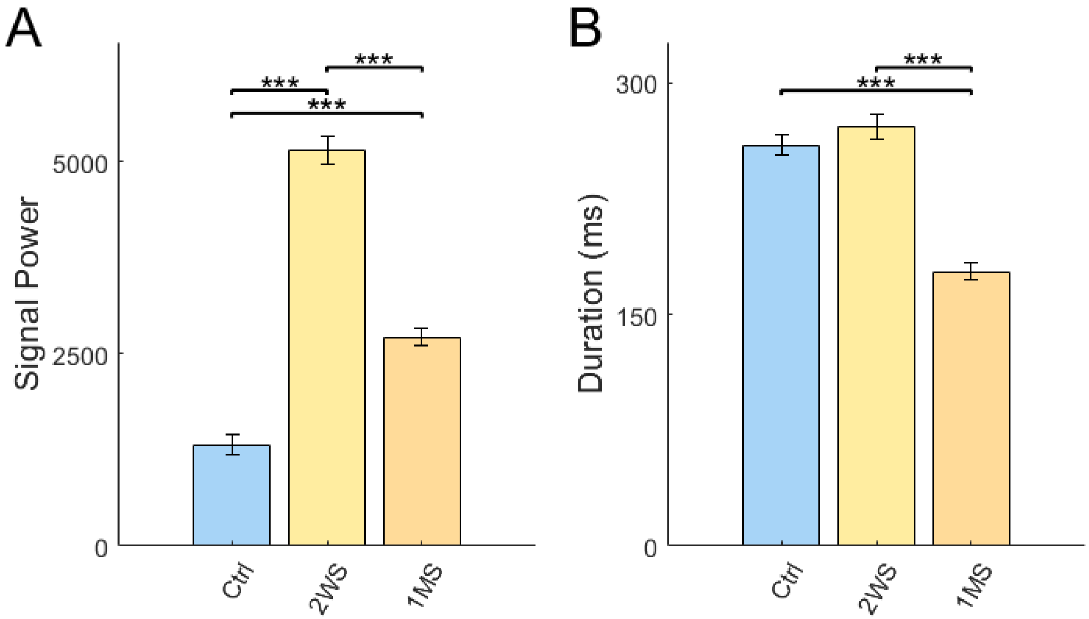


Supplementary Figure 4. Comparison of contralesional SPW-R power and duration. *Significant differences (p < 0.001, are demarked with *** ).*

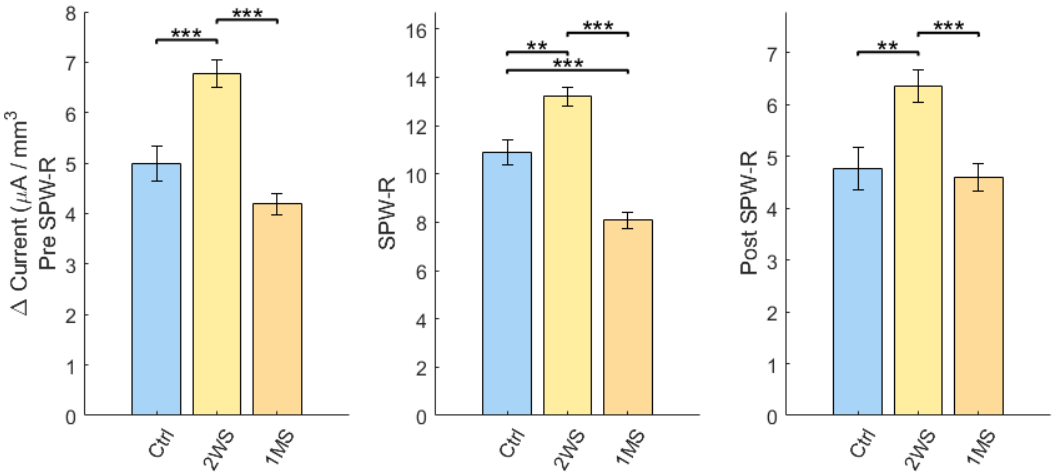


Supplementary Figure 5. Change in contralesional CSD. *Significant differences $p < 0.01$ and $p < 0.001$, are demarked with **, or *** respectively*.

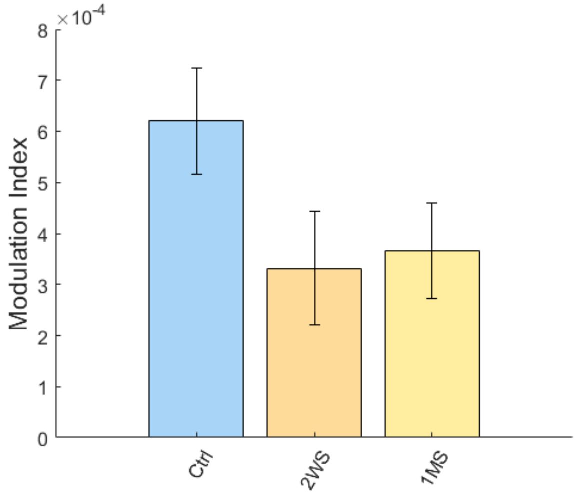


Supplementary Figure 6. Comparison of contralesional theta-gamma coupling across groups. Average modulation index within windows compared.

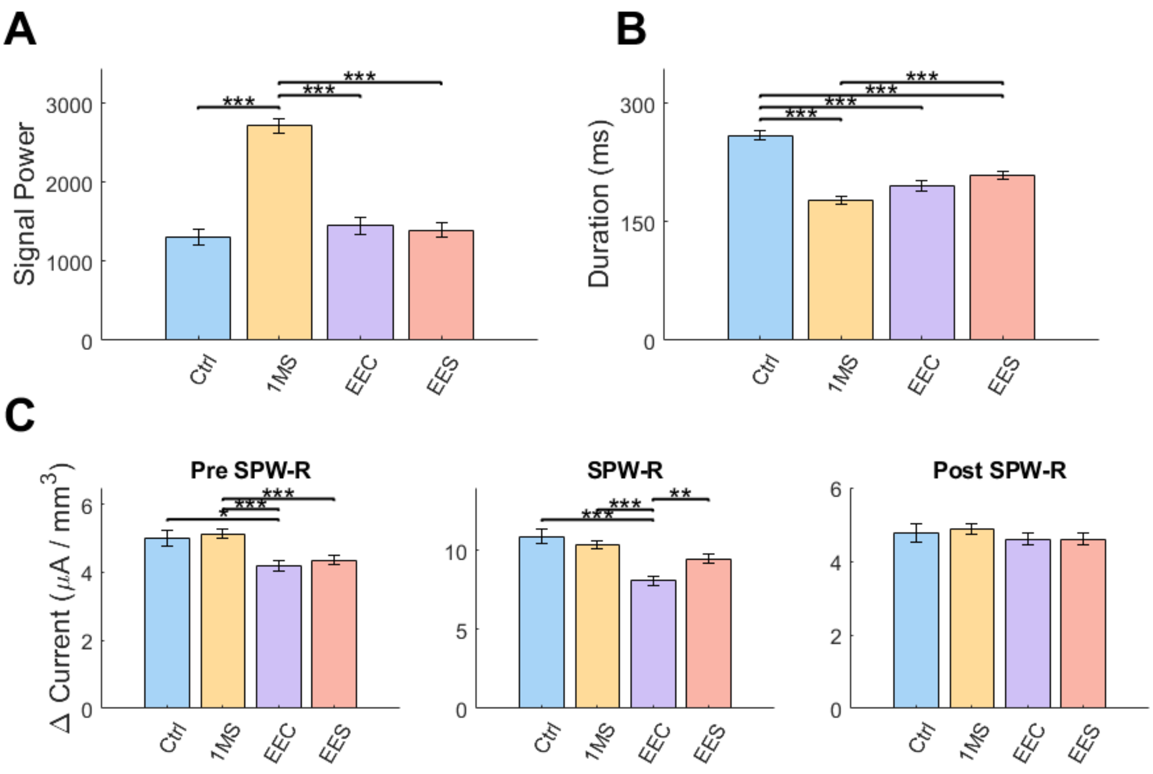


Supplementary Figure 7. Summary of the effects of EE on SPW-R biomarkers following stroke using 2-way ANOVA comparisons on contralesional hemisphere. (A) Changes in SPW-R power (B) Changes in SPW-R duration. (C) Changes in dipole amplitude surrounding SPW-R. *Significant differences: $p < 0.05$, $p < 0.01$ and $p < 0.001$, are demarked with *, **, or *** respectively*.

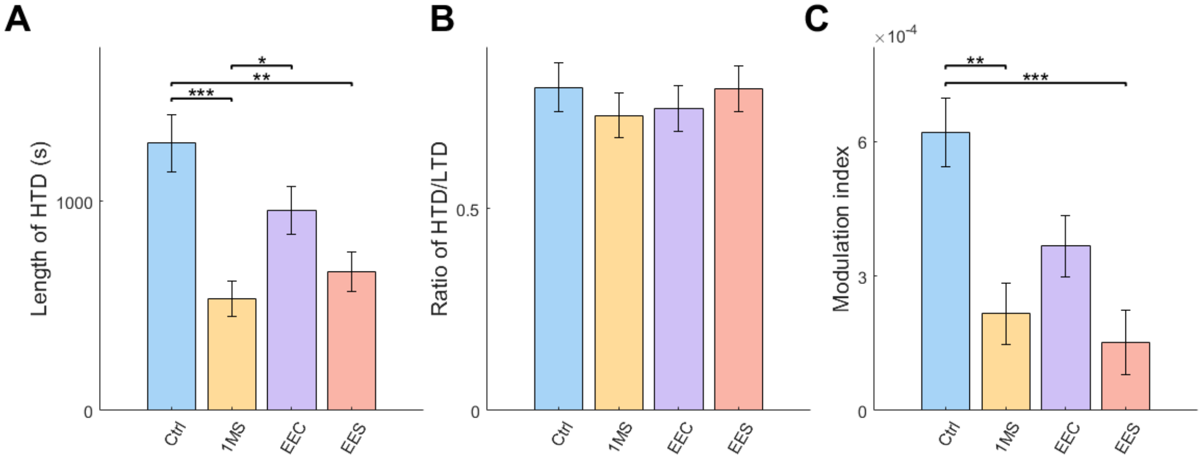


Supplementary Figure 8. Summary of the effects of EE on biomarkers disrupted following stroke using 2-way ANOVA comparisons on contralesional hemisphere. (A) Changes in average length of HTD state (B) Change in ratio of HTD/LTD. (C) Changes in theta-gamma coupling between cortex and pyramidal. *Significant differences: $p < 0.05$, $p < 0.01$ and $p < 0.001$, are demarked with *, **, or *** respectively*.
